# Hemocyte subpopulation changes in *Bithynia* snails infected with *Opisthorchis viverrini* in Thailand

**DOI:** 10.1101/536292

**Authors:** Kulwadee Suwannatrai, Apiporn T. Suwannatrai, Chalermlap Donthaisong, Patpicha Arunsan, Kavin Thinkhamrop, Pairat Tabsripair, Jariya Umka Welbat, Sirikachorn Tangkawattana, Javier Sotillo, Smarn Tesana

## Abstract

The *Bithynia* snails of *B. funiculata*, *B. siamensis siamensis* and *B. siamensis goniomphalos*, are first intermediate hosts of *Opisthorchis viverrini*. The success of parasitic infection in snails is related to the host species and efficiency of their internal defense system. Parasitic infections in snails exhibited significant variations in the number of hemocytes in hemolymph. This study investigated hemocyte counts in *Bithynia* snails for various durations of *O. viverrini* infection. The three species/subspecies snails were infected with *O. viverrini* and hemocytes were counted at post-exposed periods of 2, 4, 6, 12, 24, 48, 96 h, and 1, 4 and 8 wk to evaluate the hemocyte response to the infection. Three major hemocyte morphotypes, namely, agranulocytes, semigranulocytes and granulocytes, were observed. After infection, the differential hemocyte morphotype counts in all species/subspecies significantly increased in granulocytes at 2 h (*p* < 0.05) and decreased in semigranulocytes and agranulocytes at 2 to 4 h (*p* < 0.05). The total hemocyte counts were significant increased at 2 h after exposure in *B. s. siamensis* and *B. funiculata* (*p* < 0.05). Additionally, the number of hemocytes was reduced after exposure at 4, 6 and 12 h in *B. s. goniomphalos* and at 12, 24 and 48 h in *B. funiculata* (*p* < 0.05). From 96 h to 8 wk, the number of hemocytes was reestablished in the hemolymph, indicating that defensive cells in the host species have different mechanisms for their susceptibility or resistance to parasitic infections. Further studies on molecular functions will be done.

## 1. Introduction

The human liver fluke, *Opisthorchis viverrini*, is an important food-borne parasite in the Greater Mekong sub-region (Keiser and Utzinger, 2005; IARC, 2012). The aquatic life cycle of *O. viverrini* requires *Bithynia* snails, including *B. funiculata*, *B. s. siamensis* and *B. s.goniomphalos* as the first intermediate host; and, although the prevalence of *O. viverrini* infection is high in humans and cyprinid fish, it is very low in *Bithynia* snails in the same endemic area (Vichasri et al., 1982; Upatham et al., 1984; Brockelman et al., 1986). Furthermore, these three subspecies of snails show different levels of susceptibility/resistance to *O. viverrini* infection (Wykoff et al., 1965; Harinasuta and Harinasuta, 1984; Chanawong and Waikagul, 1991). Indeed, experimental infection studies have shown that *B. funiculata* and *B. s. siamensis* have a higher susceptibility to *O. viverrini* than *B. s.goniomphalos* (Chanawong and Waikagul, 1991). According to Prasopdee et al. (2015) a higher infection rate (61.78%) was present in early experimental infections of *B. s. goniomphalos* with *O. viverrini*, compared to later stages of infection. The differences in the susceptibility of the host and survival of the parasite in snails have been attributed to many factors such as the age, genetics, host-parasite specificity and immune system of the snail host as well as to external factors (Lie and Heyneman, 1977; Loker, 2010; Prasopdee et al., 2015).

The immune system of snails involves the innate immune response, composed by the humoral defense system, and the cell-mediated immune response (hemocytes), both operating in the processes of killing and eliminating foreign microorganisms (Foley and Cheng, 1972; van der Knaap and Meuleman, 1986; Seta et al., 1996; Yoshino et al., 2008; Comesana et al., 2012). Hemocytes have many functions, including being the main effectors of the immune system (Foley and Cheng, 1972; Seta et al., 1996; Loker, 2010; Ray et al., 2013), and present morphological and biological heterogeneities. The small cells (agranulocytes) have a relatively low immunological competence, whereas the large cells with pseudopodia (semigranulocytes and granulocytes) display enhanced immunological capacities and high phagocytic activity (Cheng, 1984; van der Knaap and Meuleman, 1986; Yoshino et al., 2008). Experimental studies have emphasized the positive effect of parasitic infections on the number of hemocytes in the hemolymph of various species (McReath et al., 1982; Bezerra et al., 1997; Ordas et al., 2000; Oliveira et al., 2010; Santos et al., 2011; Barcante et al., 2012). Furthermore, differences in the cell proportion occur often. At the onset of infection, the small cells (agranulocytes) change into granulocytes, which have a large size, numerous cytoplasmic inclusions and long pseudopodia (van der Knaap and Meuleman, 1986; van der Knaap et al., 1993; Gorbushin and Iakovleva, 2006). These granulocytes are effector cells that migrate to the infection site, thus the number of granulocytes in the hemolymph decreases (Bezerra et al., 1997; Santos et al., 2011; Barcante et al., 2012).

The aim of this study was to demonstrate the relative effect of parasitic infection by *O. viverrini* on the variations in hemocyte parameters in each species/subspecies of *Bithynia* snails. A hemocytometer was used to evaluate the total hemocyte counts and differential hemocyte morphotype counts in the snails of each species/subspecies after infection. This study provides, for the first time, a thorough characterization of the cellular response in the snail first intermediate host against *O. viverrini*, contributing to the understanding of the snail’s biology and immune responses against this devastating parasite.

## 2. Materials and Methods

### 2.1. Ethics

The protocols for animal used were approved by the Animal Ethics Committee of Khon Kaen University, based on the ethics of animal experimentation of the National Research Council of Thailand (Ethics clearance number AEKKU51/2557).

### 2.2. Snails sample preparation

All snails used in this experiment were collected from natural water bodies from the north, central and northeast regions of Thailand. We selected only medium-sized snails (shell length in cm) for this experiment: *B. funiculata* (1.0-1.2), *B. s. siamensis* (0.6-0.8) and *B. s.goniomphalos* (0.8-1.0). The snail species/subspecies was identified by using shell morphology following the available descriptions (Brandt, 1974) and verge pigmentation patterns (Jumlongpho and Tabsripair, 2012). Snails were also examined for trematode infections by using cercarial shedding and reexamined once a week for 2 months to ensure they were free from any trematode infection. Only non-trematode-infected snails were used for these studies. Snails were reared in a 50 L plastic container containing de-chlorinated tap water to a depth of 10 cm with a soil base and were provided with boiled gourd leaves and artificial snail food (Sumethanurungkul, 1970). The de-chlorinated tap water was changed every other day.

### 2.3 *O. viverrini* egg preparation

Embryonated *O. viverrini* eggs were obtained from the dung of naturally infected cats. Briefly, the dung was mixed with normal saline and strained though 4 sized meshes with 1,000-, 300-, 106- and 45-μm pores. Samples were sedimented and washed several times with normal saline in the sedimentation jar until the supernatant was clear. Subsequently, the samples were incubated at room temperature (28 ± 3 °C) for 1 wk to obtain fully developed miracidial eggs, and the normal saline was daily changed. Eggs were examined under a light microscope to determine that the miracidia were fully developed and the larvae showed active movement (Khampoosa et al., 2012).

### 2.4 Snail infection with O. viverrini eggs

Six hundred snails of each species/subspecies were used for the experiment and divided into 2 groups: infected (N = 300) and control (non-infected, N = 300). Snails (infected group) were infected with *O. viverrini* eggs by ingesting fully embryonated eggs as described previously (Prasopdee et al., 2015). Briefly, snails were exposed to 50 *O. viverrini* eggs at the optimal temperature of 28 ± 3°C in the water bath and their activity was activated under electric light. The liberation of the miracidia from the eggshell to infect the snails was determined by examining egg hatching in snail feces (as determined by opening of the operculum) (Khampoosa et al., 2012). The control group was not exposed to eggs. Finally, infection success was determined by PCR (Prasopdee et al., 2015). These snails were raised under natural conditions of dark and light and fed on synthetic snail food (Sumethanurungkul, 1970) supplemented with boiled gourd leaves. Any dead snails were removed and their number recorded every day.

### 2.5 Collection of hemolymph

Hemolymph was collected individually from each snail for analysis at 0 (control group), 2, 4, 6, 12, 24, 48 and 96 h and 1, 4 and 8 wk post-exposure by stimulation of the head-foot portion as described by Sminia (1972). Briefly, the snail shell was cleaned with 70% alcohol, the operculum was dissected, and the head-foot portion was punctured with an insulin needle. The resulting retraction of the head-foot portion deeply into the shell caused hemolymph to leak from the tissue when the operculum was removed. Then, the hemolymph was collected and transferred into a vial in an ice bath to prevent hemocyte aggregation.

### 2.6 Hemocytes subpopulation change

The total hemocyte counts (THC) and differential hemocyte morphotype counts (DHC) were determined in both infected and control groups at different time points of post-exposure. The THC was performed following the method of Barracco et al. (1993). Briefly, fresh hemolymph from individual snails (N = 20) in each experiment was immediately mixed with an anti-aggregant solution (AAS, EDTA 1.5% in phosphate buffer 100 mM, pH 7.4), stained with 0.5% neutral red solution and then fixed in 10% neutral formalin solution. Next, 10 μL of stained hemolymph was loaded onto a Neubauer hemocytometer to determine the THC under a light microscope at 400x. The THC was expressed as the number of hemocytes per milliliters. THC (cells/ml) was calculated as THC = [HC] X [DF] X [CF]/0.4, where HC was the hemocytometer count, DF was the dilution factor and CF was the conversion factor for changing cubic millimeters to milliliters. The DHC was determined under a light microscope following the method of Barracco et al. (1993). Briefly, a hemolymph sample from the same individual snails as for the THC (N = 20) was stained with neutral red and placed on a slide. The cell morphology and granularity of the hemocytes were examined using permanent staining. A total of 200 hemocytes were counted per slide to ascertain the hemocyte morphotypes, and then the relative percentages were calculated.

### 2.7 Statistical analyses

Data referring to the THC and percentages of DHC counts at different time points are presented as means and standard deviation. Comparisons between the control and infected snails at the different time points in each species/subspecies were analyzed using Student’s t-test, with *p* < 0.05 being significant.

## 3. Results

### 3.1 Experimental infection rate

The survival rates of experimentally infected snails until the end of experiment in *B. funiculata*, *B. s. siamensis* and *B. s. goniomphalos* were 93.33%, 91.67%, and 96.67% (280, 275 and 290 out of 300 snails), respectively. In infected groups, egg hatching increased over the course of the infection, with more than 50% egg hatching as seen at 24 h after exposure (Fig 1A). Infection was detected by PCR in *B. funiculata* and *B. s. goniomphalos* at 4 h until 8 wk post-exposure, (and negative in control group, 0 h) while, in *B. s. siamensis*, it was detected from 2 h until 8 wk. Infection rate was very low (0-35%) in early infection (2–6 h) while it reached 35-80% at 12 h-1 wk following a decrease at 4-8 wk. The average infection rate of infected snails was 27% in *B. funiculata*; 38% in *B. s. siamensis*; and 39% in *B. s. goniomphalos*. The highest infection rate was observed in *B. s. goniomphalos* at 24 h post-exposure (80% by PCR) (Fig. 1).

**Fig. 1.**
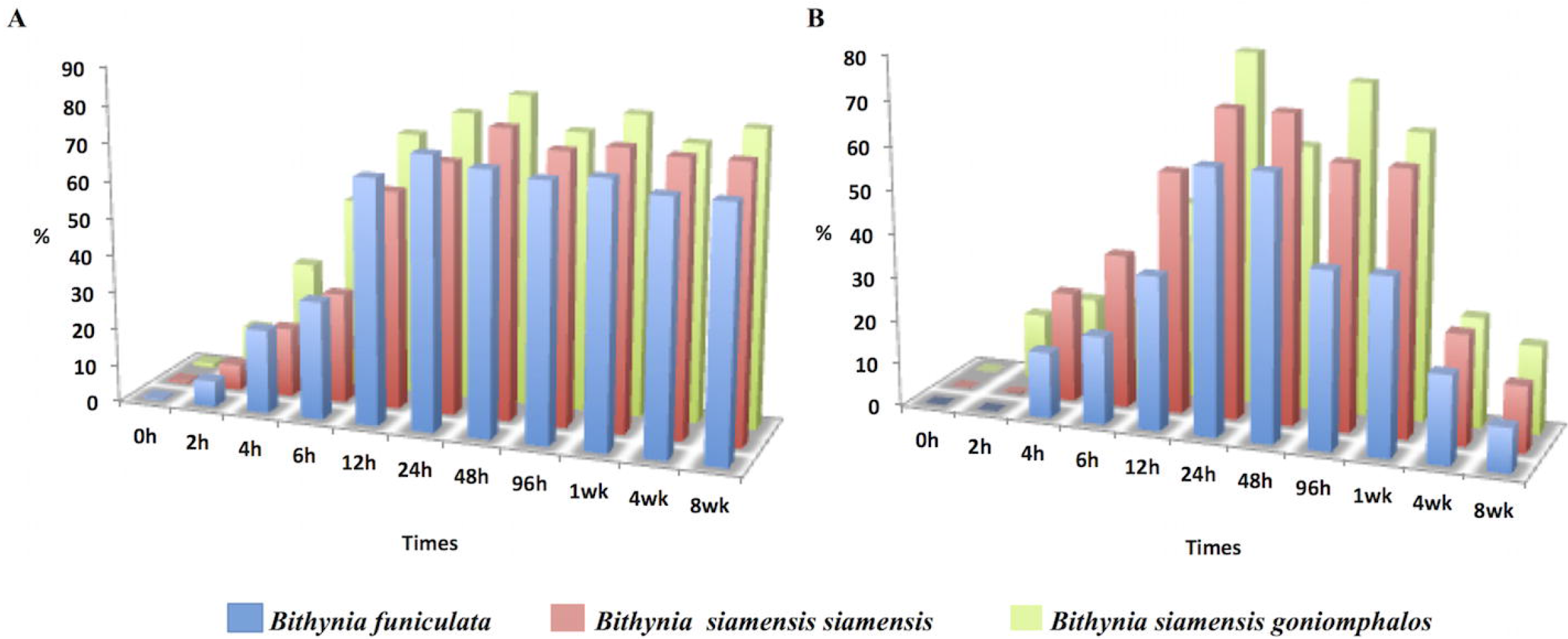
Egg hatching rates (A) and infection rates by PCR analyses (B), in *Bithynia funiculata*, *Bithynia siamensis siamensis* and *Bithynia siamensis goniomphalos* after exposure to *O. viverrini* eggs

### 3.2 Hemocyte morphotype of B. funiculata, B. s. siamensis and B. s. goniomphalos

The characterization of hemocyte morphotypes was performed following previous studies (McCormick-Ray and Howard (1991), Accorsi et al. (2013) and Ray et al. (2013)) by neutral red staining according to size and shape of cells and nuclei, nucleus/cytoplasm (N/C) ratio, presence/absence of cytoplasmic granule and the presence of pseudopodia. Hemocytes in the control groups of *B. funiculata*, *B. s. siamensis* and *B. s. goniomphalos* revealed 3 major hemocyte morphotypes of agranulocytes, semigranulocytes and granulocytes (Fig. 2).

**Fig. 2.**
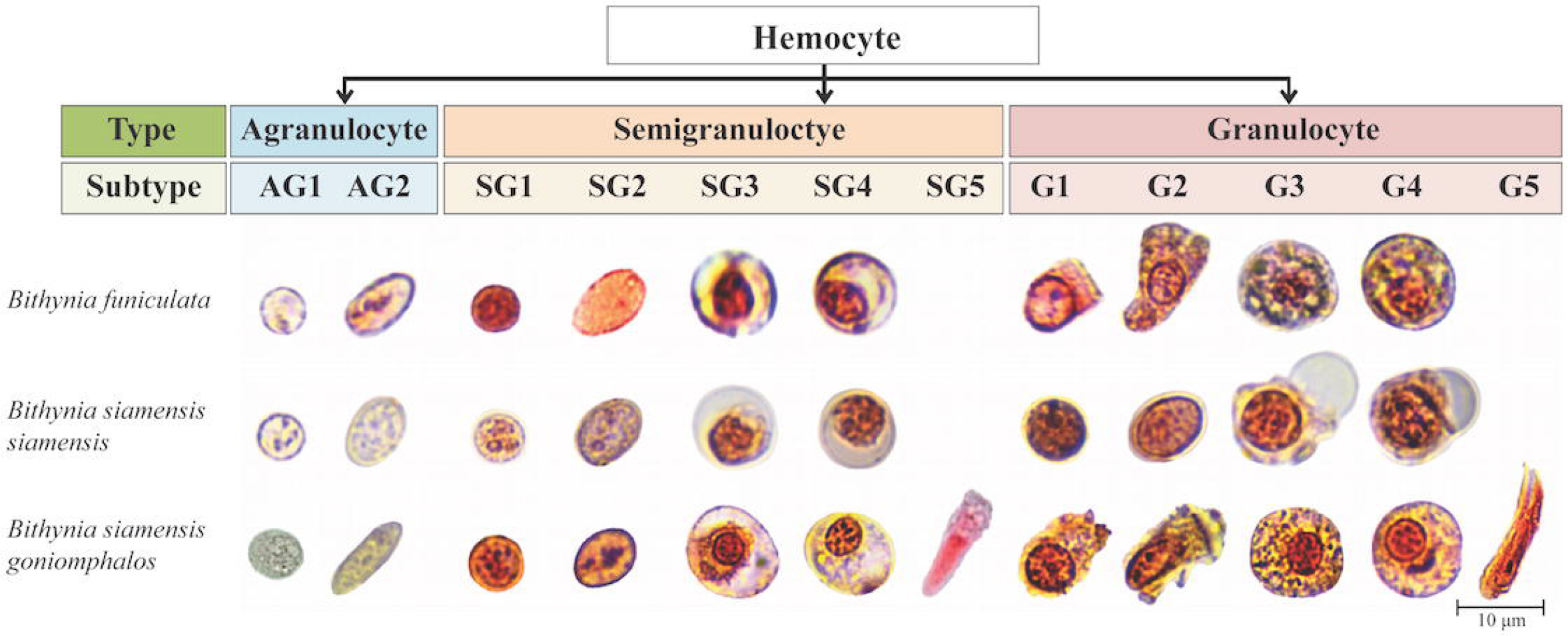
Hemocyte types from *Bithynia* snails stained with neutral red and observed under light microscope. Scale bar=10 μm

Agranulocytes (AG) had no granules, were blast-like cells and light red stained. The majority of agranulocytes from the *Bithynia* snails were small in size, had large nuclei surrounded by a thin rim of cytoplasm with a high nucleus/cytoplasm (N/C) ratio and occasionally showed short pseudopodia spreading. They included 2 subtypes of round shape (AGI), and oval shape (AGII) (Fig. 2, Table 1).

**Table 1.**
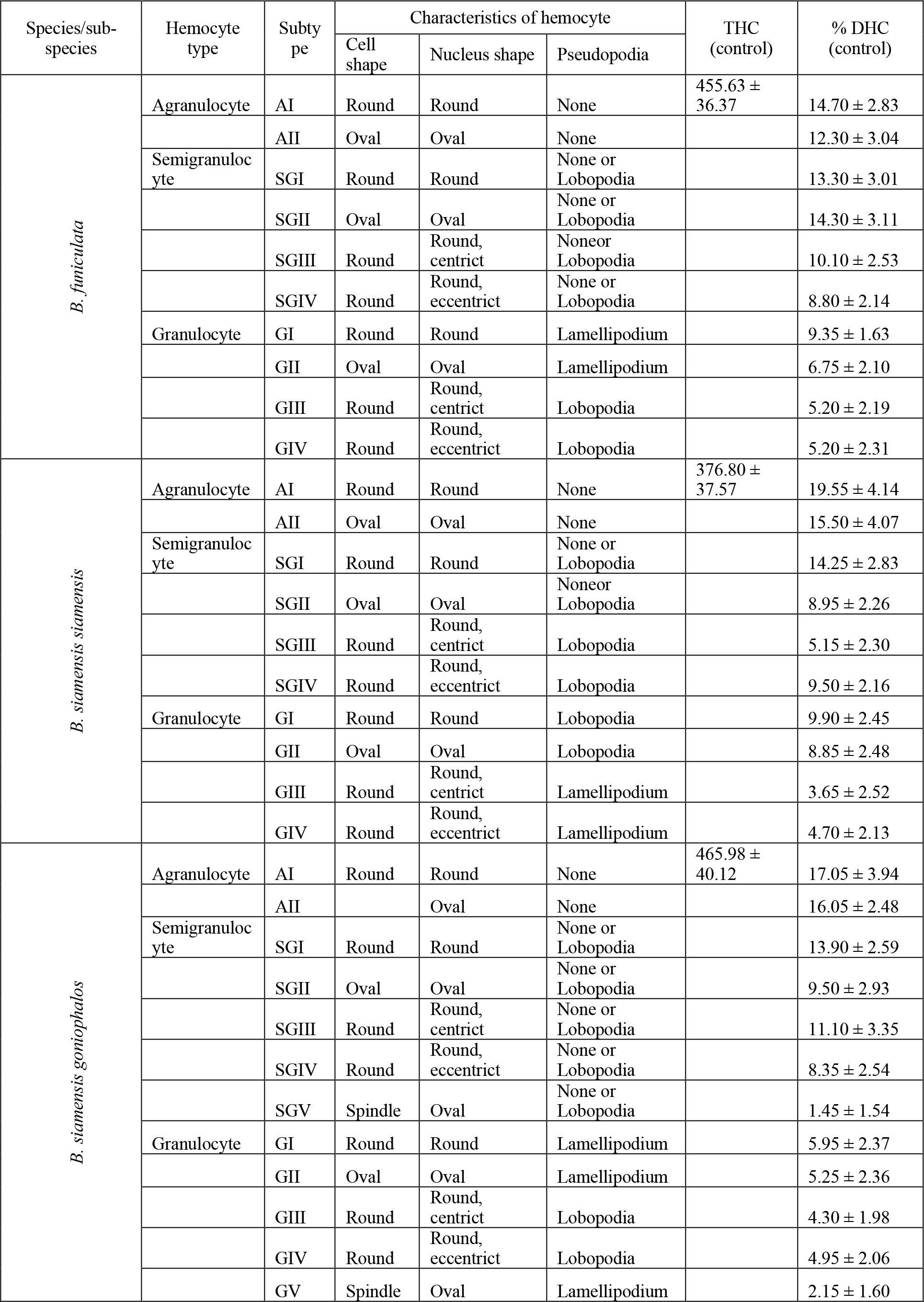
Summary of the hemocytes characteristics in *Bithynia funiculata*, *Bithynia siamensis siamensis* and *Bithynia siamensis goniomphalos*

Semigranulocytes (hyalinocytes) constituted the most abundant population of hemocytes in the hemolymph. Their cytoplasm contained few red-colored granules, finger-liked pseudopodia (lobopodia) and low N/C ratio. Finger-like pseudopodia (lobopodia) could be present in the small size semigranulocytes (SG), which could be divided into 2 subtypes based on shape: round (SGI) and oval (SGII). The large size semigranulocytes with lobopodial pseudopodia could be classified into 3 different subtypes based on shape: round shape with centric nuclei (SGIII) or eccentric nuclei (SGIV) and semigranulocyte subtype V (SGV), which was spindle-shaped with lamellipodial pseudopodia and was only found in *B. s. goniomphalos*.

Granulocytes (G) were larger with abundant red granules in their cytoplasm and pseudopodia. The small-sized granulocytes could be classified into 2 subtypes based on shape: round (GI) and oval (GII). The large-sized granulocytes could also be classified into 2 subtypes: round shape with centric (GIII) or eccentric (GIV) nuclei. Granulocyte subtype V (GV) was spindle shaped and was only found in *B. s. goniomphalos* (Fig. 2, Table 1).

### 3.3 Total hemocyte count (THC)

The range of total hemocyte counts (mean±SD) in the non-infected groups (controls) of *B. funiculata*, *B. s. siamensis* and *B. s. goniomphalos* were 392.25-512.25 (455.62 ± 36.37), 315.50-438.00 (376.80 ± 37.56) and 370.50-561.75 (465.98 ± 40.12) × 104 cells/ml, respectively. In the infected groups, the number of circulating hemocytes was significantly increased in *B. s. siamensis* (*p* < 0. 05) and *B. funiculata* (*p* < 0. 05) at 2 h after infection. However, the increase in THC from *B. s. goniomphalos* was not significant. The THC gradually decreased from 4 h to 12 h after infection in *B. s. goniomphalos* and from 12 h to 48 h in *B. funiculata* compared with the non-infected groups (*p* < 0.05). Interestingly, during the same period of time, the THC was temporarily increased in *B. s. siamensis* (2 h to 24 h, *p* < 0. 05) compared with the non-infected group. From 96 h post-exposure with *O. viverrini*, the total number of hemocytes plateaued until 8 wk in all species/subspecies (Fig. 3).

**Fig. 3.**
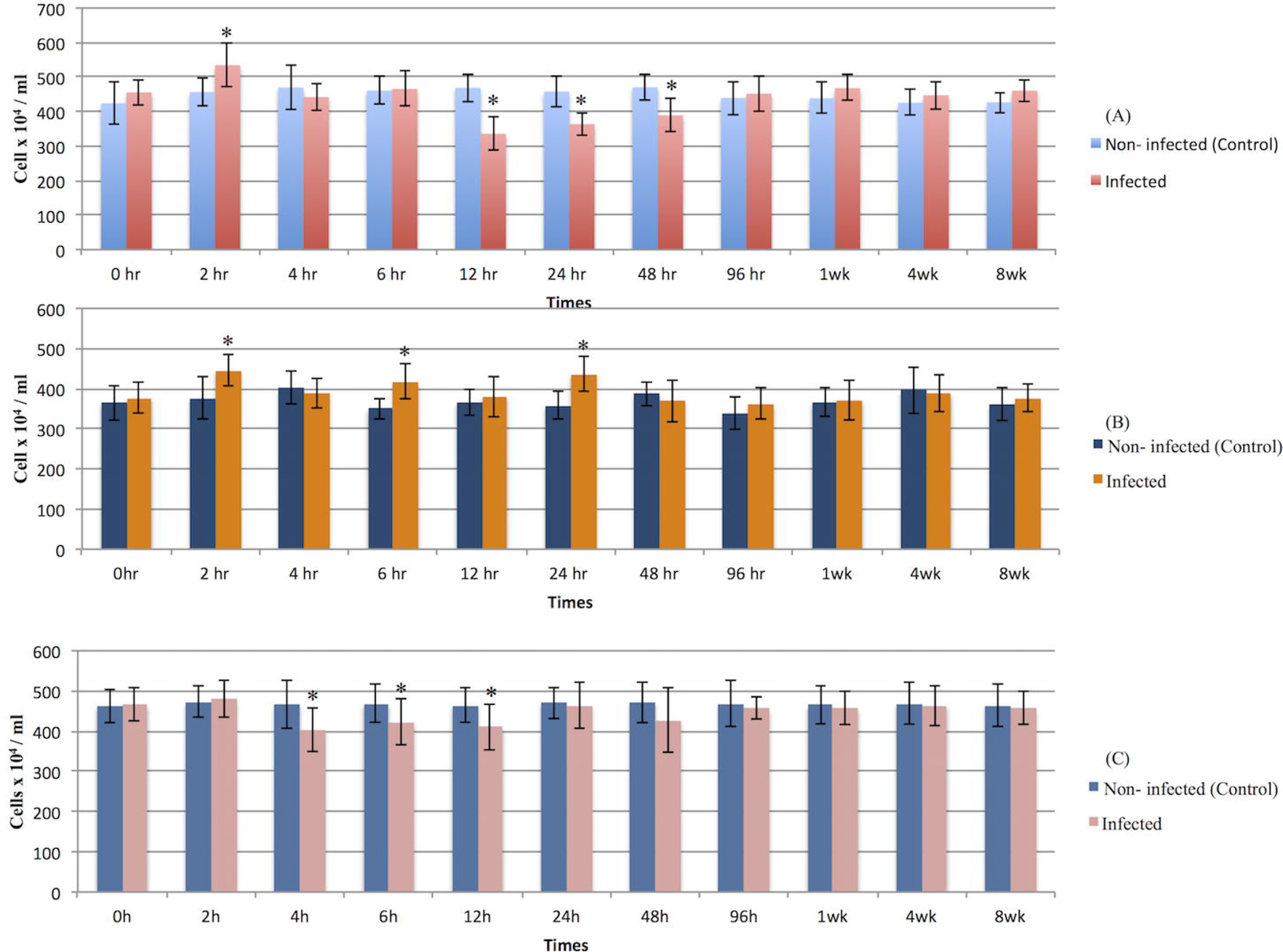
Total hemocyte count (THC) in the hemolymph of *Bithynia funiculata* (A), *Bithynia siamensis siamensis* (B) and *Bithynia siamensis goniomphalos* (C), after exposure to *O. viverrini* eggs at 0 (control group), 2, 4, 6, 12, 24, 48, and 96 h and 1, 4 and 8 wk post-exposure.

### 3.4 Differential hemocyte morphotype counts (DHC)

The hemocyte morphotypes of *B. funiculata*, *B. s. siamensis* and *B. s. goniomphalos* were divided into 3 major groups of agranulocytes, semigranulocytes and granulocytes and subdivided into 12 subtypes. In non-infected groups (control groups), the majority of circulating cells were semigranulocytes, followed by agranulocytes, with few granulocytes. After infection, there was an increase in granulocytes and a decrease in agranulocytes and semigranulocytes compared with the control groups (Fig. 4).

**Fig. 4.**
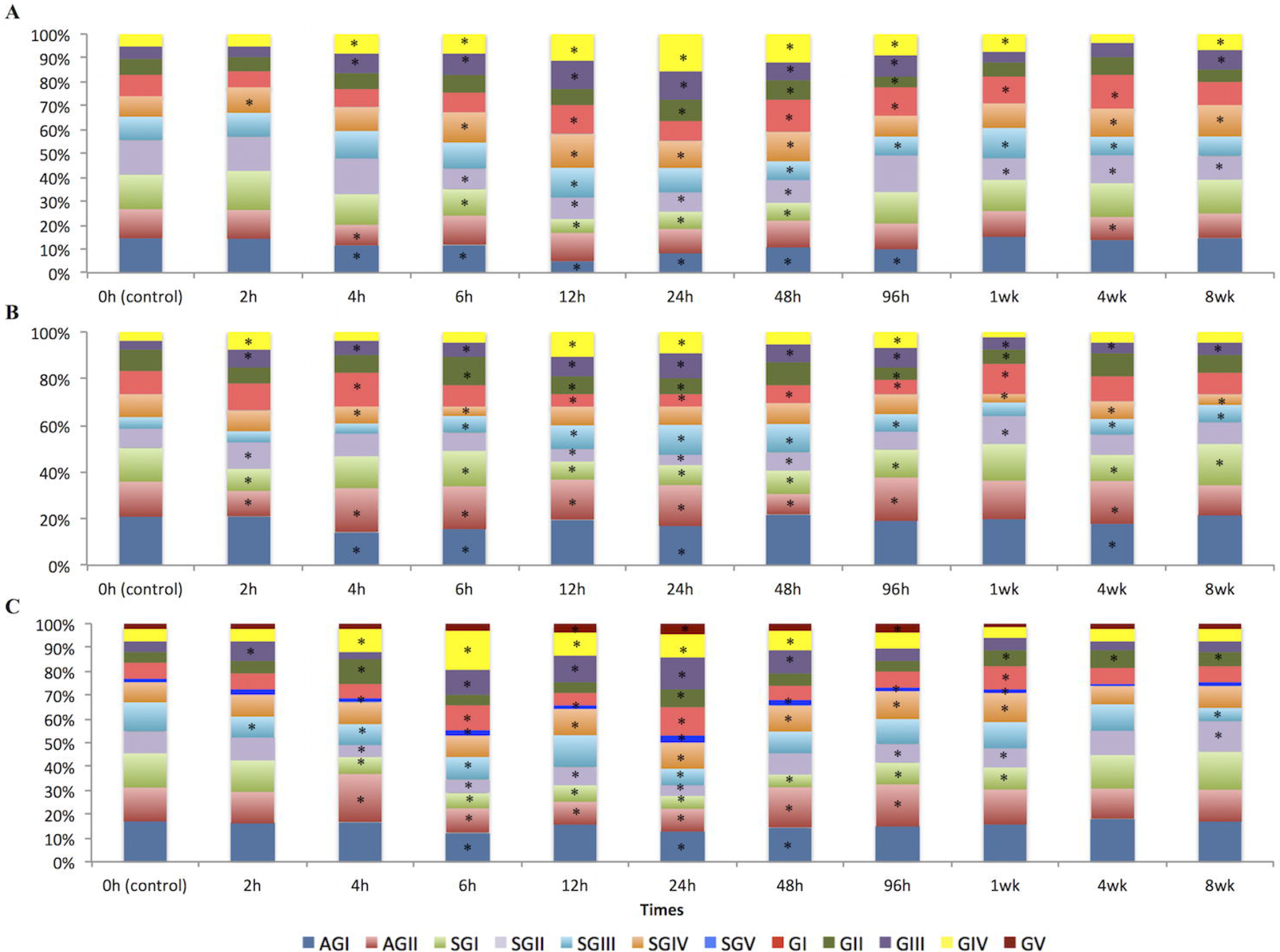
Differential hemocyte count (DHC) in the hemolymph of *Bithynia funiculata* (A), *Bithynia siamensis siamensis* (B) and *Bithynia siamensis goniomphalos* (C) after exposure to *O. viverrini* eggs at 0 (control group), 2, 4, 6, 12, 24, 48, and 96 h and 1, 4 and 8 wk post-exposure. Mean number of each circulating hemocyte subpopulation is shown. Agranulocyte subtype I (AGI), agranulocyte subtype II (AII), semigranulocyte subtype I (SGI), semigranulocyte subtype II (SGII), semigranulocyte subtype III (SGIII), semigranulocyte subtype IV (SGIV), semigranulocyte subtype V (SGV), granulocyte subtype I (GI), granulocyte subtype II (GII), granulocyte subtype III (GIII), granulocyte subtype IV (GIV), granulocyte subtype V (GV). * represents significant differences (p < 0.05) compared to control groups.

In infected *B. funiculata*, the percentage of AGI and AGII was significantly decreased at 4 h compared to non-infected snails while the percentage of SGIII, GIII and GIV was significantly increased (p<0.05) compared with non-infected snails. At 6 to 24 h, the percentage of AGI, SGI and SGII was significantly decreased while the percentage of SGIII, SGIV, GIII and GIV was increased (p<0.05) in infected snails (Fig. 4A). At 48 to 96 h, the percentage of AGI, SGIII was significantly decreased while the percentage of GI, GIII and GIV was significantly increased (p<0.05) (Fig. 4A). At 1 wk, the percentage of SGII was significantly decreased while the percentage of SGIII, GI and GIV was significantly increased (p<0.05) (Fig. 4A). Finally, at 4 to 8 wk, the percentage of SGII and SGIII was significantly decreased while the percentage of SGIV and GIV was significantly increased (p<0.05) (Fig. 4A).

In infected *B. s. siamensis*, at 2 h, the percentage of AGII and SGI was significantly decreased compared with non-infected snails while the percentage of SGII, GIII and GIV was significantly increased (p<0.05) (Fig. 4B). At 4 to 6 h, the percentage of AGI was significantly decreased while the percentage of GIII was increased (p<0.05) (Fig. 4B). At 12 to 96 h, the percentage of SGI, SGII, GI and GII was decreased while the percentage of AGII, GIII and GIV was increased (Fig. 4B). At 1 wk, the percentage of SGIV was significantly decreased while the percentage of SGII, GI and GIII was significantly increased (p<0.05) (Fig. 4B). Finally, at 4 to 8 wk, the percentage of SGI and SGIV was significantly decreased while the percentage of SGIII and GIII was significantly increased (p<0.05) (Fig. 4A).

In infected *B. s. goniomphalos*, at 2 h, the percentage of SGIII was significantly decreased compared with non-infected snail while the percentage of GIII was significantly increased (p<0.05) (Fig. 4C). At 4 to 48 h, the percentage of AGI, AGII, SGI, SGII and SGIII was decreased while the percentage of SGIV, SGV, GI, GII, GIII, GIV and GV was increased (p<0.05) (Fig. 4C). At 96 h to 1 wk, the percentage of SGI and SGII was significantly decreased while the percentage of SGIV and SGV was significantly increased (p<0.05) (Fig. 4C). Finally, at 4-8 wk, the percentage of GII was significantly increased (p<0.05) (Fig. 4C).

## 4. Discussion

Early studies classified snail hemocytes into 2 main types: agranulocytes (or hyalinocytes) and granulocytes (Feng et al., 1971; Sminia, 1972; Cheng, 1984). Recently, a third type of hemocytes was described hemocytes, allowing researchers to classify hemocytes into 3 main types: agranulocytes, semigranulocytes and granulocytes (Gorbushin and Iakovleva, 2006; Ray et al., 2013). In the present study, we were able to classify hemocyte morphotypes from three different species intermediate hosts of *O. viverrini* into 3 main types with 12 subtypes, including 2 subtypes of agranulocytes (blast-cell liked), 5 subtype of semigranulocytes (hyalinocytes) and 5 subtypes of granulocytes. Surprisingly, only one species, *B. s. goniomphalos*, presented spindle-shaped hemocytes (SGV and GV). This classification suggests that the morphology, staining with specific dyes, sizes and relative granularity of the cytoplasm were extensions of previous hemocyte classifications (McCormick-Ray and Howard, 1991; Accorsi et al., 2013; Ray et al., 2013). Agranulocytes of *Bithynia* snails had small cell sizes and large nuclei. These cells had a poorly defined role in defense and reactions to antigens. Preliminary observations had suggested that agranulocytes were not involved in phagocytosis and the agglutination of foreign material (Cheng, 1975; van der Knaap and Meuleman, 1986; van der Knaap et al., 1993). In contrast, semigranulocytes and granulocytes had larger cell sizes with pseudopodia, present morphological variations and display enhanced immunological capacities. However, granulocytes had more abundant granules in the cytoplasm than semigranulocytes. Therefor, granulocytes could be classified as the main factors of the snail’s immune response against against parasitic infection (Sminia, 1972; Cheng, 1984; Yoshino et al., 2008).

The life cycle of *O. viverrini* requires *Bithynia* snails as the first intermediate host. The snail becomes infected by ingestion of parasite eggs, which hatch the miracidium in their digestive tract, penetrate the host tissue and invade host defense mechanisms to establish within their hosts (Wykoff et al., 1965; Harinasuta and Harinasuta, 1984). Prasopdee et al. (2015) reported that, after *Bithynia* snail ingested parasite eggs or food, the snail depleted the ingested food from gut within 20 to 340 min (average 161.10 ± 63.43 min). As a result, parasite eggs were retained in the snail’s digestive tract allowing for false positive results when infection rate was determined only by PCR. To avoid this problem we provided them with boiled gourd leaves to deplete the ingested parasite egg before collecting the hemolymph. According to previous studies, *B. funiculata* and *B. s. siamensis* have higher susceptibility to *O. viverrini* than *B. s. goniomphalos* (Chanawong and Waikagul, 1991).

However, the present study has not shown different infection rates in the different *Bithynia* spp. In addition, the highest infection rate was observed at 24 h-1wk post-exposure and proportionally decreased in the long term of infection (4-8 wk p.i.), which is similar to previous reports (Prasopdee et al., 2015). In addition the PCR analyses were supported by egg- hatching observations, with lower egg-hatching in early infections. The lower number of egg hatching was found in control group (*O. viverrini* eggs incubated in the mixture of snail feces), suggesting that egg hatching depends on several factors, especially the factors from snail such as enzymes (Khampoosa et al., 2012).

In mollusks, hemocyte production occurs in the amebocyte product organ, in the hematopoietic organ and in peripheral vascular locations over time (Lie et al., 1975; Rondelaud and Barthe, 1981; Sminia, 1981). Snails have an open circulatory system with hemocytes moving in the hemolymph. These cells engage in diapedesis, with the hemocytes passing into tissue spaces and ingesting foreign materials (Lie et al., 1975; Sminia, 1981). After infection, the internal defense system (IDS) of snails were capable of activating the cellular and humoral system for the elimination of the parasite. This activation could be a direct stimulation by the pathogen. For example, excretory/secretory products and extracts of the parasite stimulate the enlargement and increased mitosis of the amoebocyte-producing organ in *B. glabrata*, contributing to changes in the number of hemocytes in hemolymph (Noda, 1992). The number of hemocytes in hemolymph has been established in several mollusks, varying greatly from species to species and depending on physiological condition of the snails such as infection with pathogens (Sminia, 1981; Barcante et al., 2012; Ray et al., 2013); however, no report exists yet on *Bithynia* snails. Evaluation of the number of hemocytes in the hemolymph is usually expressed as the THC, while morphological differences among hemocytes is expressed as the DHC (Barracco et al., 1993; Donaghy et al., 2009; Comesana et al., 2012). For a better understanding of the snail-parasite interactions, the present study has demonstrated the relative effect of *O. viverrini* infection on the hemocyte population in each species/subspecies of *Bithynia* snails at different time points. Marked differences in the hemocyte population were found among *Bithynia* snails infected with *O. viverrini* and this finding was supported by PCR and egg-hatching of infected snails. At 2 h after exposure, THC in the hemolymph samples of all snail species increased. After that (4 h to 96 h), the THC gradually decreased in *B. s. goniomphalos* and *B. funiculata*, while temporary significantly increased in *B. s. siamensis*. Then, at 1 wk until 8 wk, the total hemocytes plateaued. This correlates with previous experimental parasite infections in *B. glabrata*, where the THC was significantly reduced at 5 h after infection with *S. mansoni* (Bezerra et al., 1997; Santos et al., 2011) and reduced at 4 h up to 72 h after infection with *A. vasorum* (Barcante et al., 2012). After that, the number of hemocytes was reestablished at 10-15 days after infection (Seta et al., 1996; Bezerra et al., 1997; Oliveira et al., 2010). This trend is possibly a result of the snail’s immune response, which is initiated with the first parasite infection and continued until parasites were eliminated, hence showing variation in the number of hemocytes in the hemolymph of infected snails.

The DHC results showed a small alteration of the hemocyte subtypes was present in all snails at the beginning of the infection period (0-2 h), with semigranulocytes representing the majority of hemocyte types followed by agranulocytes and granulocytes. The highest alteration in the DHC was observed at 4-96 h post-exposure. In *B. funiculata*, the percentage of small agranulocytes and semigranulocytes (AGI, AGII, SGI and SGII) decreased while the percentage of large semigranulocytes and granulocytes (SGIII, SGIV, GI, GII, GIII and GIV) significantly increased. In *B. s. siamensis*, the percentage of all small hemocytes (AGI, SGI, GI and GII) was significantly decreased while all large hemocytes (SGIII, SGIV, GIII and GIV) were increased. In *B. s. goniomphalos*, the percentage of all small hemocytes (AGI, AGII, SGI and SGII) decreased but the percentage of large semigranulocytes (SGII and SGIV) and all granulocytes (GI, GII, GIII, GIV and GV) significantly increased. In the later stages of infection (1-8 wk), the proportion of semigranulocytes and granulocytes was still increased. According to previous studies granulocytes are effector cells, while the small cells (agranulocytes) are considered as an early stage of the larger cells (Foley and Cheng, 1972; van der Knaap and Loker, 1990; Gorbushin and Iakovleva, 2006). However, the relative decrease in one cell type will cause an apparent increase in the others. For example following exposure of *B. s. goniomphalos* to *O. viverrini*, the percentage of AGI, AGII, SGI and SGII (small cells) were significantly decreased while the percentage of SGIV, SGV, GI, GII, GIII, GIV and GV (large cells) was increased at 24 h (high infection rate). This observation suggests that that AGI is an earlier stage of SGI, then maturing to SGIII and SGIV and finally differentiated to GIII or GIV. Similarly, AGII is earlier stage of SGII, then maturing to SGV and developing to GII or GV, with semigranulocytes acting as immediate precursors of granulocytes. The decrease in small agranulocytes and semigranulocytes and increase in large semigranulocytes and granulocytes in the hemolymph after infection could be a result of the penetration of the parasite in the snail’s tissue and the release of excretory/secretory products, which could directly stimulate the defense system of snails, resulting in an increase hemocyte production in amoebocyte-producing organ of snail leading to change proportion of hemocyte in the hemolymph in early infection. Then, small cells could act as a precursor of the larger cells that can differentiate into mature effector cells leading to a decrease proportion of small cells but to an increase proportion of large cells (Noda and loker, 1989; Foley and Cheng, 1972; van der Knaap and Loker, 1990; Gorbushin and Iakovleva, 2006; Oliveira et al., 2010; Ray et al., 2013). Moreover, during the course of infection, the presence of hemocytes in the hemolymph and tissue might be interpreted as a result of a response from the snail’s immune system, providing more effector cells to the infection sites, resulting in a decreased number of granulocyte in the hemolymph.

This study suggested that parasitic infections emphasize a positive effect on the number of hemocytes in the hemolymph and each snail species presents different IDS to protect them from parasitic infections. Furthermore, more studies are needed to clarify the role of hemocytes and better understand the immune response of *Bithynia* snails to *O. viverrini* infection.

## Acknowledgements

This work was supported by the Development and Promotion of Science and Technology Talents Project (DPST) scholarship from the Institute for the Promotion of Teaching Science and Technology, Royal Thai Government, Thailand.

